# COCOA: A Framework for Fine-scale Mapping Cell-type-specific Chromatin Compartmentalization Using Epigenomic Information

**DOI:** 10.1101/2024.05.11.593669

**Authors:** Kai Li, Ping Zhang, Jinsheng Xu, Zi Wen, Junying Zhang, Zhike Zi, Li Li

## Abstract

Chromatin compartmentalization and epigenomic modification are crucial factors in cell differentiation and diseases development. However, mapping precise chromatin compartmental patterns across multiple cell types requires Hi-C or Micro-C data at high sequencing depth. Exploring the systematic relationship between epigenomic modifications and compartmental patterns remains a challenge. To address these issues, we present COCOA, a deep neural network framework that uses convolution and attention mechanisms to infer reliable fine-scale chromatin compartment patterns from six representative histone modification signals. COCOA achieves this by extracting 1-D track features through bi-directional feature reconstruction after resolution-specific binning epigenomic signals. These track features are then cross-fused with contact features using an attention mechanism. Subsequently, the contact features are transformed into chromatin compartment patterns through residual feature reduction. COCOA demonstrates accurate inference of chromatin compartmentalization at a fine-scale resolution and exhibits stable performance on test sets. In addition, we explored the impact of histone modifications on the chromatin compartmentalization through *in silico* epigenomic perturbation experiments. When using 1kb resolution high-depth experimental data, obscure compartments are observed, whereas COCOA can generate clear and detailed compartmental patterns. Finally, we demonstrated that COCOA enables cell-type-specific prediction of unrevealed chromatin compartment patterns in various biological processes. Thus, COCOA is an effective tool for gaining chromatin compartmentalization insights from epigenomics in a wide range of biological scenarios.

## Introduction

The three-dimensional (3D) architecture of chromatin is essential for gene expression during cell differentiation and disease development [1,2]. Recent advances in next generation sequencing have led to the development of several chromosome conformation capture techniques, such as Hi-C, Micro-C, and Pore-C [3–5], enabling the exploration of multiscale chromatin structural elements including chromatin compartment [3,6], topological associating domains (TADs) [7,8], loops [6], stripes [9] and microcompartments [10]. These techniques have revealed that the chromatin can be segregated into A compartment and B compartment [3,11]. The A compartments are generally active in the genome, whereas the B compartments are mostly transcriptional repressive. These chromatin compartments are closely related to the mechanisms underlying various key biological processes [12,13].

To identify chromatin compartments, sequencing data is usually processed into contact maps and distance effects are eliminated using normalization methods. The normalized contact map is then used to calculate the correlation matrix (CM), which is subjected to principal component analysis (PCA). The sign of the first principal component (PC1) corresponds to the compartment state [3]. Most analyses related to chromatin compartments rely on CM and PC1 [14–16]. While the CM is commonly available and of high-quality at mega-base scale, it becomes noisy for resolution finer than 25kb, failed to show clear plaid patterns due to its sparseness. Recent studies have suggested the associations of the fine-scale resolution chromatin compartments with other structural elements [17,18], histone modifications and chromatin accessibility [19]. However, the available chromatin compartment data does not match the scale of the epigenomic data, making the connection between epigenomics and chromatin compartmentalization a challenge. Furthermore, due to technical limitations and sequencing costs [20], experimentally mapping high-resolution chromatin compartment is both expensive and labour-intensive. Therefore, there is unmet need for the development of a computational method to obtain the fine-scale CM across multiple cell lines.

In the past decade, deep learning [21] has emerged as a widely used tool in computational 3D genomes. These applications include various tasks such as TAD boundary recognition [22,23], chromatin loop detection [24,25], chromatin interaction data enhancement [26,27], interaction matrix generation [28–30] and single cell Hi-C imputation [31,32]. While several methods explore contact map generation and enhancement, they lack cell type specificity. For example, HiC-Reg [33] used fourteen epigenomic signals from five cell lines to predict short-range chromatin interactions using random forests. Akita [29] and Orca [34] adopted the convolutional neural network to predict contact maps from DNA sequences. However, these methods are not capable directly inferring contact maps across different cell types. Recently, two proposed methods, C.Origami [35] and Epiphany [30], address this limitation by utilizing the histone modifications and chromatin accessibility data. C.Origami predicts short-range interactions by integrating CTCF, chromatin accessibility, and DNA sequence information through a neural network containing the attention and convolutional modules. Epiphany uses multiple epigenomic signals to generate short-range chromatin contact maps. However, these methods have their own limitations in terms of chromatin compartmentation and method generalization. Firstly, these existing methods concentrate on the prediction of short-range interactions (TADs and loops) while ignoring long-range interactions (compartments). Additionally, the relationship between compartmentalization and histone modification is still unresolved. Furthermore, these models require inputs in fixed bin sizes, limiting scalability and preventing across-resolution predictions.

To address these limitations, we introduce COCOA (mapping **c**hr**o**matin **co**mpartmentalization with epigenomic inform**a**tion), a method that predict the cell-type-specific CM using six types of accessible epigenomic modification signals. COCOA adopts bidirectional feature reconstruction and cross attention fusion for bi-directional reconstruction and fusion of epigenomics data. Subsequently, residual feature reduction is applied to map the fused results into CM. COCOA is specifically designed to generate chromatin compartmentalization, and the predicted CM can be directly used to determine compartment status. We evaluated the performance of COCOA using multiple metrics (MSE, MAE, GenomeDISCO, PCC, SSIM and PSNR). The results demonstrate that COCOA accurately generates significant and biologically meaningful CM. Furthermore, we conducted *in silico* perturbation experiments to investigate the influence of histone modifications on compartments. Additionally, we tested the generalization performance of COCOA by making model predictions with resolution-specific and cell-type-specific data. The results show that COCOA enables robust performance at various resolutions from diverse cell lines, providing insights into the patterns of chromatin compartments in immune and disease tissues.

## Method

### Hi-C and Micro-C data sources and pre-processing

We collected publicly available processed Hi-C and Micro-C data of different cell lines from the 4DN database [36]. Intra-chromosomal contact maps were computed from these data for model training and testing (see details in Table S1). Depending on the specific task, the intra-chromosomal contact maps is computed at different resolutions using the cooler package [37]. To eliminate the distance effect in the contact maps, we applied the observed-expected normalization method [3]. Finally, these normalized contact maps were converted into CMs, which clearly depict plaid pattern of chromatin compartmentalization.

### ChIP-seq data sources and pre-processing

Histone modification signals (H3K9me3, H3K27ac, H3K4me1, H3K27me3, H3K4me3, H3K36me3) from the ChIP-seq [38] data for all cells were retrieved from ENCODE project [39] (see details in Table S2). The ChIP-seq data were binned to specific resolutions using the pyBigWig package (Figure 1A). After binning, a log(x + 1) transformation and min-max normalization were performed on the data. Finally, the processed data were combined into an epigenomic signal matrix.

**Figure 1.**
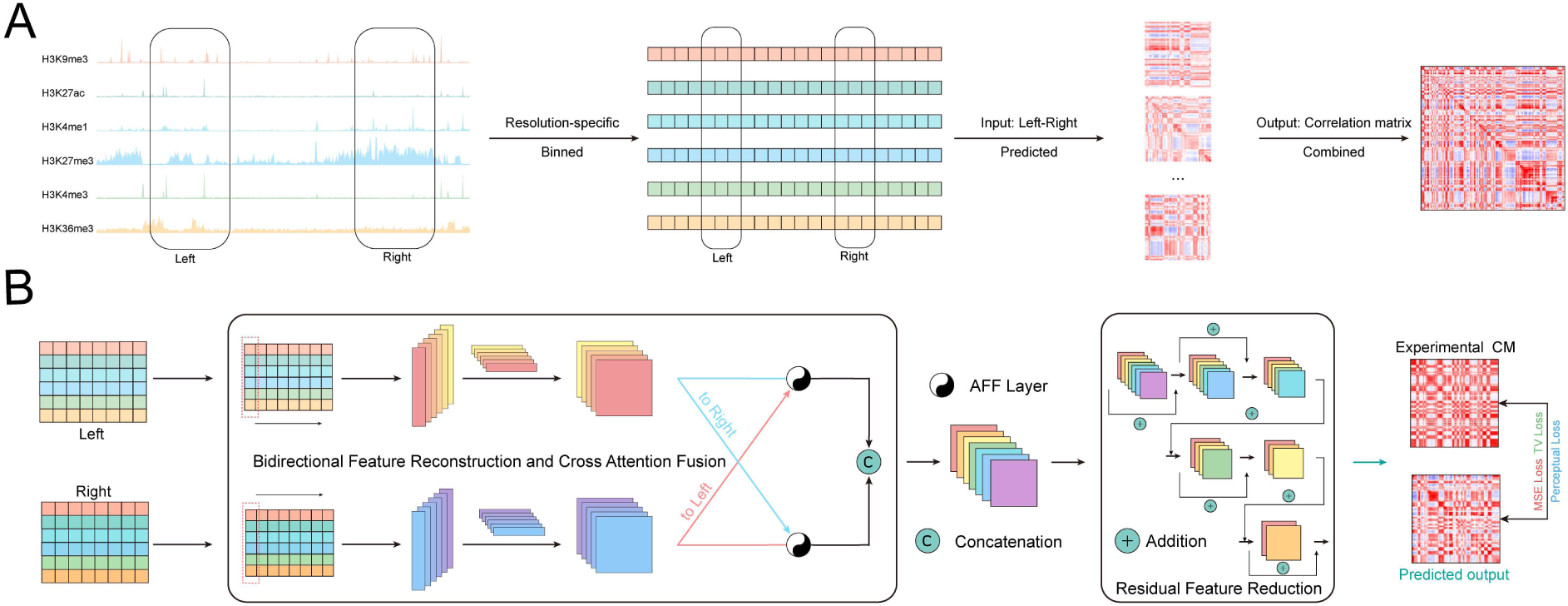
COCOA pipeline and architecture. **(A)** COCOA pipeline: The integration of six accessible epigenomic signals by resolution-specific binning serves as inputs to predict the correlation matrices. **(B)** COCOA architecture: COCOA extracts 1D track features from each input (bidirectional feature reconstruction module) and then combines these features with spatial contact features (cross attention fusion module). The contact features are further processed by the residual feature reduction module to obtain the final prediction result. Parameters are updated using a backpropagation algorithm with mixed loss functions. (Refer to Method for detailed information)

### Dividing matrices

The pre-processing step generates two matrices: a symmetric correlation matrix (CM) with dimensions n×n ( CM_n×n_) and an epigenomic matrix (EM) with dimensions m×n ( EM_m×n_). Each value CM_ij_ in the CM_n×n_ represents the correlation strength between genomic segment i and j. Values greater than one indicate that the two genomic segments have the same interaction mode, while less than one indicate the opposite interaction mode, providing information about the status of chromatin compartments. Each value EM_ij_ in the EM_m×n_ represents the signal strength of genomic segment j on epigenomic track i.

To better preserve the plaid pattern of chromatin compartmentalization and adapt to the inputs of neural network, we implemented the following processing scheme. First, the CM_n×n_ was divided into sub-matrices of k × k size (SCM_k×k_), and the EM_m×n_ were divided into two sub-matrices of m × k size (SLEM_m×k_ and SREM_m×k_). We started at the diagonal position in the top-left corner of CM_n×n_ and moved horizontally, dividing it into SCM_k×k_. Simultaneously, we divided the two corresponding groups of genomic loci from EM into SLEM_m×k_ and SREM_m×k_. After completing the horizontal dividing, we moved the current position diagonally by k positions. This process was repeated until the entire CM_n×n_ could no longer be divided. Due to computational resource constraints, we sampled the SCM_k×k_ in groups to minimize the size of the training datasets (SLEM_m×k_ and SREM_m×k_ were synchronized to minimize). Finally, these data were saved separately for further modelling.

### Combining predicted sub-matrices

The COCOA model takes the SLEM_m×k_ and SREM_m×k_ for each chromosome division as inputs. It then outputs a series of predicted correlation sub-matrices. These sub-matrices sequentially covered a square matrix (PCM_n×n_) with the same number of columns as the EM_m×n_. The specific coordinates for covering each predicted correlation sub-matrices are determined by the corresponding inputs (SLEM_m×k_ and SREM_m×k_). Finally, the complete PCM_n×n_ is generated and saved for further biological analyses.

### COCOA architecture

The COCOA model consists of three main components: bidirectional feature reconstruction, cross attention fusion, and residual feature reduction (Figure 1B), which are described in the following sections.

### Bidirectional feature reconstruction

The bidirectional feature reconstruction module consists of two matrix reconstruction layers (MR layers). The construction of these MR layers is inspired by our previous work on chromatin interaction data enhancement [40]. Each MR layer consists of two parts: an aggregation convolution layer with a filter size of N × 1 and a linear reconstruction layer. The output of each MR layer is computed as follows:

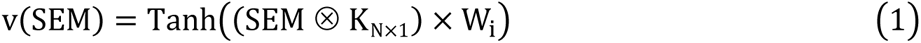

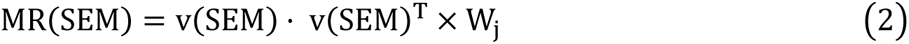

where ⊗ denotes the convolution operation, K_N×l_ represents convolution kernel (N × 1), Tanh is the activation function [41], × denotes Hadamard product and · denotes dot product. SEM represents SLEM_m×k_ or SREM_m×k_ generated through pre-processing. v^T^ represents the transposition of the vector v. W_i_ and W_j_ are learnable weight matrices, respectively. The MR layer aggregates multiple ChIP-seq track signals from different genomic loci into a 1D vector. This vector is in turn reconstructed into a low-ranking epigenomic track features using learnable weight matrices. In summary, this module obtains bi-directional epigenomic track features by reconstructing the SLEM_m×k_ and SREM_m×k_.

### Cross attention fusion

Next, the COCOA model employs the cross attention fusion module to fuse bi-directional epigenomic track features. The cross attention fusion module mainly contains two attention feature fusion layers (AFF layers) [42]. Each AFF layer has three parts: global feature extraction, local feature extraction and attention fusion. The results of cross attention fusion are defined as follows:

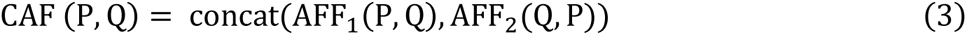

where P and Q represent bi-directional epigenomic track features, respectively. concat denotes stacking two outputs in the same dimension. AFF refers to an attention-based uniform and general neural network layer for feature fusion proposed by Dai et al. [42]. The cross attention fusion module transforms the epigenomic track features in the other direction into potential attention weights to reinforce the epigenomic track features in the current direction. By interleaving attention fusion and concatenation, a set of fused contact feature maps is obtained as inputs for the next module.

### Residual feature reduction

The residual feature reduction module consists of a series of residual blocks, each containing several residual layers. Following the approach described in previous work [43], each residual layer is composed of convolutional layers with different convolution kernels, BN layers [44], and activation functions. The computation of each layer is defined as follows:

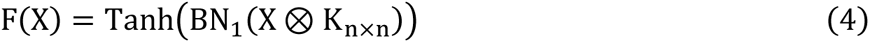

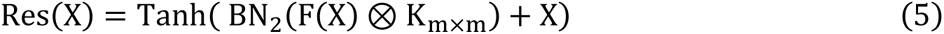

where K denotes the convolution kernels of different size, Tanh is activation function and BN is the BatchNorm layer. X represents the fused contact feature maps for the first layer and the output of the current layer serving as the input for the next layer. The residual feature reduction module decreases the channels of the contact features from the previous module, level by level. Throughout this process, the residual layer continuously filters to retain important information from the previous layer, aggregating it with the output of current layer. Finally, the predicted correlation sub-matrix is obtained from the last layer of residual feature reduction module.

### Loss function

The COCOA model can be seen as a function F with a parameter set θ, which maps each group input SLEM_i,m×k_ and SREM_i,m×k_ to the predicted correlation sub-matrix PSCM_i,k×k_ (i.e., PSCM_i,k×k_ = F(SLEM_i,m×k_, SREM_i,m×k_ ∶ θ)). The objective of the training is to find a set of θ^∗^ to enable PSCM_i,k×k_ similar to the ground truth SCM_i,k×k_. Therefore, COCOA initially uses the mean square error (MSE) loss to minimize the pairwise error of genomic range k × k between PSCM and SCM. This loss can be described as:

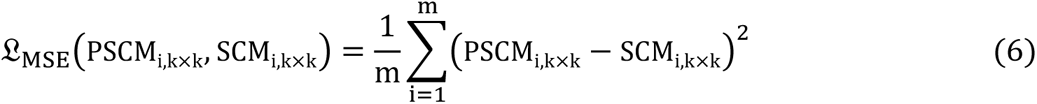

Subsequently, COCOA incorporates a perceptual loss based on the VGG network [45] to restore structural information of the correlation matrix. Furthermore, we added the total variation (TV) loss [46], which effectively smooths noise in computer vision, as a regularization term to suppress the noise of the PSCM_k×k_. These losses can be described as:

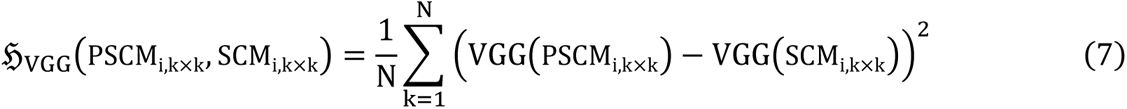

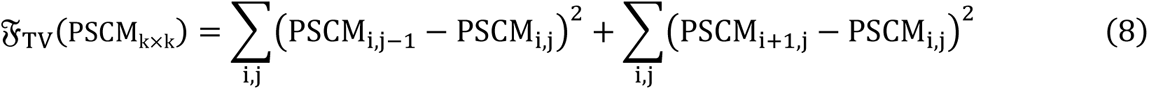

Finally, the training objective can be represented as:

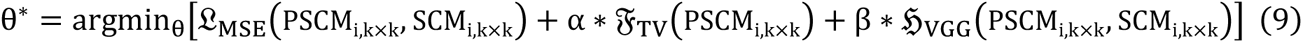

where α,β are scaling weights that range from 0 to 1.

### COCOA training and hyper-parameter exploration

Before model training, we pre-processed each chromosome of the HFFc6 Micro-C data [47] and corresponding ChIP-seq data. Chromosome 1, 3, 5, 7, 9, 11, 13, 15, 17 and 19 were used as training sets, while chromosomes 18, 20, 21 and 22 were utilized for hyper-parameter tuning. The remaining chromosomes were allocated for performance evaluation.

The COCOA model was implemented on Python 3.7 with PyTorch1.12 [48]. We trained the model with a batch size of 16 for 120 epochs, using the Adam optimizer [49] with an initial learning rate of 5e-4 (lr_init_ = 5e − 4). The details on model training and hyper-parameters are provided in Note S1.

### Model evaluation

We started the evaluation process by making predictions on independent test sets using the best-trained model. The predicted correlation sub-matrices were then combined to form the intra-chromatin correlation matrix. The experimental 25 kb chromatin interaction correlation matrices were considered as the ground truth. During the evaluation, we used the principal component analysis (PCA) provided by the sklearn package [50] to calculate the first principal component (PC1) of the two correlation matrices. PC1 is generally considered to represent the A/B compartment information. Additionally, we discretized PC1 to obtain the chromatin compartment state, which was saved separately.

To assess the model performance, we used several indictors including Mean Square Error (MSE, see Equation (6)), Mean Absolute Error (MAE), Structure Similarity Index (SSIM) (assessing the similarity of two correlation matrices) and Peak Signal to Noise Ratio (PSNR) (measuring the quality score of the correlation matrices) [51]. MAE, SSIM and PSNR are defined by the following equations:

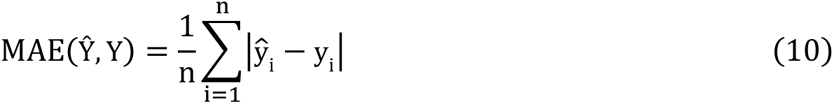

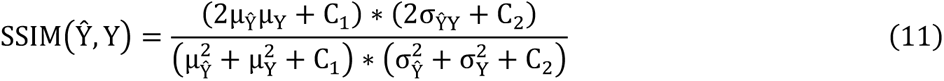

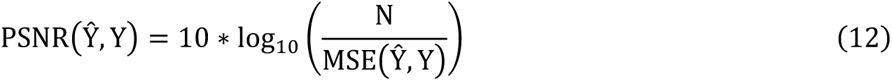

where Ŷ denotes the predicted correlation matrices and Y represents the real correlation matrices. Furthermore, considering the chromatin compartmentalization information of the correlation matrices, we evaluated their reproducibility using multiple Pearson Correlation Coefficient (PCC) and GenomeDISCO score [52].

## Results

### Overview of COCOA

In this study, we proposed COCOA as a method for accurately predicting cell-type-specific chromatin compartment patterns at fine-scale resolution by integrating epigenomic modification signals. COCOA only requires six epigenomic track signals as inputs, which are accessible for most tissues and cell lines in the ENCODE database [53]. The targets of COCOA are defined as the correlation matrices (CM) of OE-normalized contact maps, allowing for the maximum retention of chromatin compartment pattern information. The COCOA framework connects these inputs and targets through binning, prediction and combination operations (Figure 1A). Notably, in the binning process, we utilized the resolution-specific binning approach (i.e., *Bin_epi_* = *Bin_corr_*) instead of a single bin from each genome site approach (i.e., *Bin_epi_* = *Bin_corr_* ∗ *resolution*). This choice greatly improves the practicality of COCOA.

We trained COCOA on Micro-C data of HFFc6 along with corresponding ChIP-seq data (Table S1 and S2) using backpropagation algorithm. Specifically, COCOA first utilizes the bidirectional feature reconstruction module to calculate the 1D track features separately from two inputs. This step captures the intrinsic association present in the original epigenomic data in each direction (Figure 1B and Method). Subsequently, the cross attention fusion module integrates these 1D track feature maps with space contact features based on crossed attention mechanisms (see details in Method). Lastly, the residual feature reduction module decodes these contact features to generate predicted results, which are then combined into a complete CM (Refer to the Combining predicted sub-matrices section of Method for detailed information). In addition, a composite loss function is employed to minimize the distance between the predicted targets and the ground truth.

### COCOA accurately predicts chromatin compartmentalization pattern

To assess the performance of COCOA, we applied the trained COCOA model to randomly selected epigenomic data in test sets (Chr 12, Chr 14 and Chr 16) to generate predicted CM. We considered the CM calculated from the Micro-C data and its PC1 as the experimental CM, which can be regarded as the ground truth for comparison. Heat maps in Figure 2A and Figure S1A compare typical genomic regions using the predicted and the experimental CM. The results demonstrate that the predicted CM generally exhibit the correct chromatin compartmentalization pattern. Furthermore, COCOA shows outstanding generative capacity in capturing subtle chromatin compartments. Notably, the predicted CM show more pronounced interactions in dissimilar chromatin compartment blocks (blue blocks) compared with the experimental CM, while exhibiting partial over-reinforcement in identical chromatin compartment blocks (red blocks). We also computed the PC1 of the predicted CM and the experimental CM using the sklearn package [50]. Subsequently, the CMs were sorted based on the size of their respective PC1 values (Figure 2B and Figure S1B). The results indicate that modularity phenomenon of the predicted CMs resemble the modularity patterns observed in the experimental CMs. In addition, the predicted CMs successfully capture the white band regions present in the experimental CM (Figure S1B).

**Figure 2.**
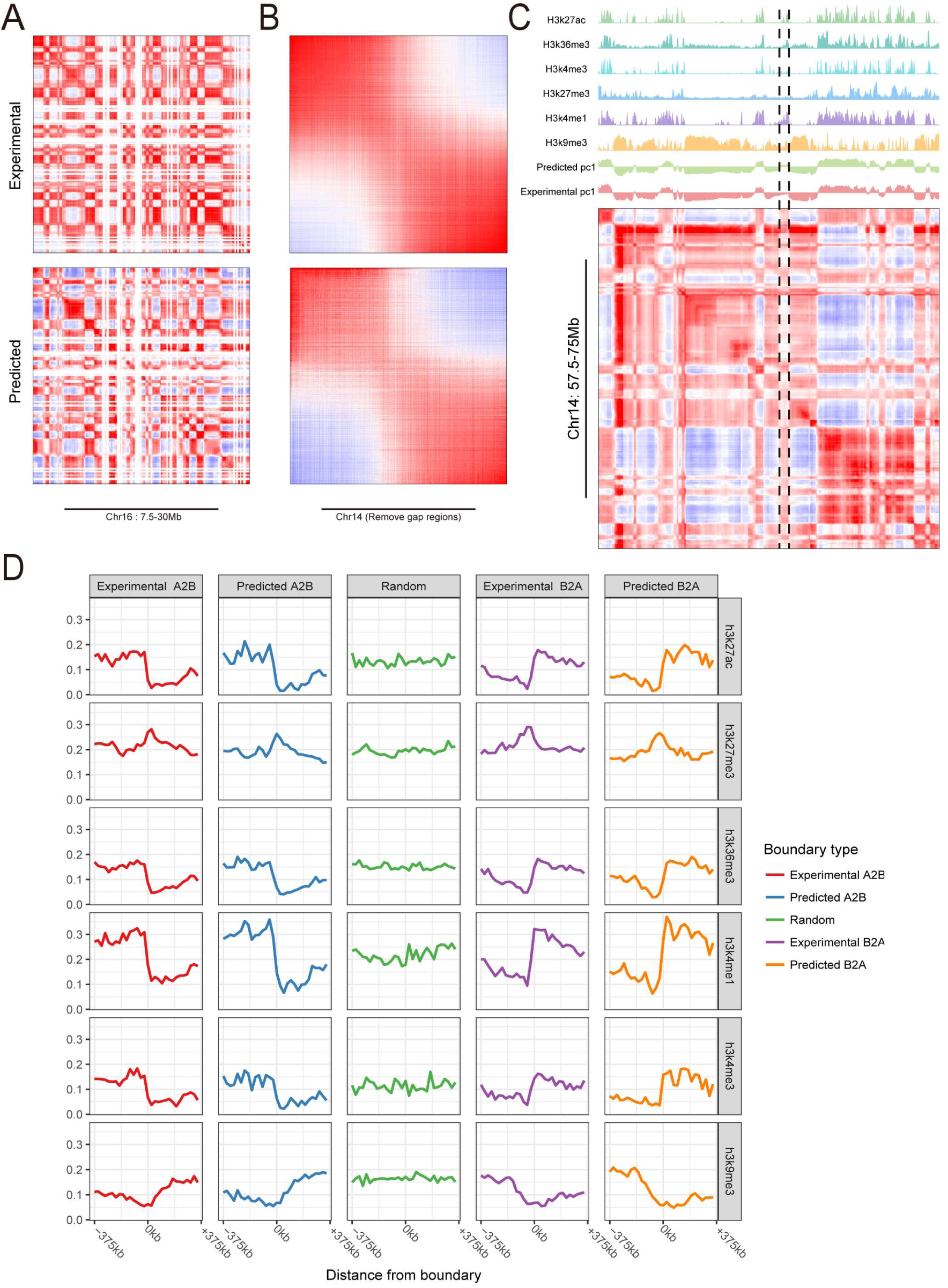
COCOA accurately predicts significant compartment patterns from epigenomic data. **(A)** Representative region illustrating predicted and experimental CM on test chromosomes. **(B)** Heatmaps of the experimental CM and the predicted CM, sorted according to their respective PC1 sizes. The predicted CM demonstrates consistency with the compartment patterns observed in experimental CM. **(C)** The predicted CM exhibits patters that align precisely with the waveform of histone modification signal. Within the region marked by the black dashed line, COCOA is able to correct the pattern misclassified by the experiment. **(D)** Analysis of the shifts in modification signals within 375 kb neighbourhoods surrounding compartment boundaries in both predicted and experimental CMs.

To establish the biological significance of COCOA model predictions, we generate plots that illustrate the predicted CMs alongside the epigenomic signal tracks and the PC1 tracks. Figure S1C reveals that the predicted CM accurately show plaid patterns of chromatin compartments, with each block of the plaid corresponding to a signal peaks in the epigenomic data tracks. The PC1 values from the tracks of the experimental CM and the predicted CM also align precisely with these results. Importantly, COCOA can infer chromatin compartments that are consistent with the underlying epigenomic data but are not captured in the experimental CM (indicated by the black dotted lines in Figure 2C). Moreover, we analysed shifts in six epigenomic modification signals at compartment boundaries and randomly selected genomic loci, as done in previous studies [54]. Notably, we observed the consistent significant shifts in epigenomic modification signals within 375 kb neighbourhoods around A/B compartment boundaries in both predicted and experimental CMs. These shifts were obviously different from randomly selected genomic loci (Figure 2D and Figure S1D). It is worth noting that shifts of partial epigenomic modification signals of the predicted CM generated by COCOA outperformed the experimental CM in capturing some compartment boundaries (e.g.: A2B boundary of H3K4me1 shown in Figure 2D).

### Genome-wide performance evaluation of COCOA

The performance of COCOA was quantitatively analysed on genome-wide test sets. We calculated the MSE, MAE, PNSR, and SSIM scores to evaluate the robustness of error, signal-to-noise ratios and the structural similarity of COCOA on test sets. COCOA achieves competitive error and similarity scores on the test sets, exhibiting only minimal fluctuation with variations in the quality of the input data and the chromosome size. This stability indicates that COCOA performs consistently across different prediction scenarios (Figure 3C, left panel and Table S3). In addition, we adopted GenomeDISCO scores, designed to assess the reproducibility of contact maps, to validate the biological significance of the predicted CM. As shown in Figure 3C (right panel), COCOA achieved high reproducibility between the predicted CM and the experimental CM.

**Figure 3.**
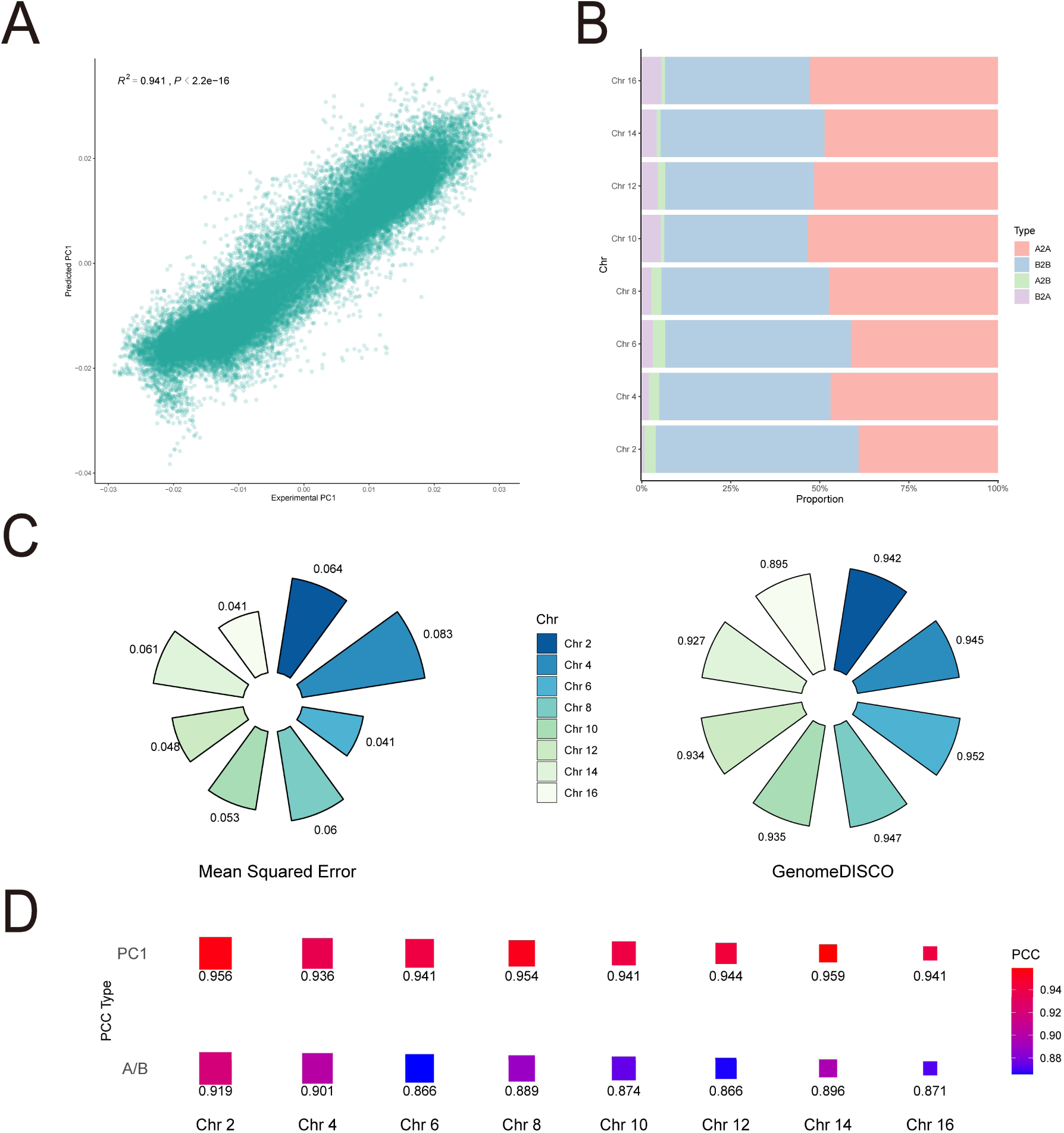
Performance evaluation of COCOA in multiple metrics. **(A)** Scatterplot showing the high correlation between the PC1 of the predicted CM and the PC1 of the ground truth on test sets. **(B)** Proportion of compartment pattern matching between the predicted CM and the experimental CM. The red and blue bars represent the proportion of compartments that overlap between the predicted CM and experimental CM. The green and purple bars indicate the proportion of compartments that differ between the predicted CM and experimental CM. **(C)** The MSE and genomeDISCO scores for COCOA on the test chromosomes. **(D)** Correlation coefficient between the predicted CM and experimental CM. The “PC1” row represents the correlation of the first principal component of the CM and the “A/B” row represents the correlation after binarization of PC1.

As the CM contains abundant information regarding chromatin compartmentalization, we preformed correlation evaluations at the CM, PC1 values and the compartment states levels. Figure 3A shows a scatter plot of the PC1 values between the predicted CM and the experimental CM across the test sets (R^2^ = 0.941, p < 2.2e − 16). In Figure 3D, the correlation coefficients of the PC1 values for each predicted and experimental CM are higher than 0.9, with the same trend observed for the A/B chromatin state correlation coefficients. Furthermore, we observed that misclassification rates of A/B compartments were independent of chromosome length and all remained below 10% (Figure 3B). To evaluate COCOA’s performance in inferring deep chromatin compartmentalization information, we calculated the mean correlation coefficients for each column between the predicted CM and the ground truth. The results show that the predicted CM achieved high correlation scores when compared to the experimental CM (Figure S2).

### COCOA predicts chromatin compartmentalization changes to epigenomic perturbations

After confirming the accuracy of COCOA in inferring chromatin compartmentalization from epigenomic data, we used COCOA to perform *in silico* epigenomic perturbation experiments and assessed the impact of epigenomic signals on chromatin compartment patterns.

In the single epigenomic marker perturbation (one-perturbation) experiments, we generated perturbed epigenomic data by setting one selected epigenomic signal to its minimum value while keeping other data unchanged. Subsequently, we predicted the corresponding CMs for the perturbed epigenomic data and compared them to the experimental CM for the unperturbed data. The predicted results from one-perturbation experiments indicated that alerting H3K9me3 signal significantly influenced chromatin architecture, causing a substantial number of compartment B-to-A switches (Figure 4A). On the other hand, the perturbation of H3K4me1 signal led to a small proportion of compartment A-to-B switches. Perturbing other epigenomic signals (e.g., H3K27ac, H3K27me3, H3K4me1, and H3K4me3) had no significant effect on chromatin compartment patterns (Figure 4A and Figure S3A).

**Figure 4.**
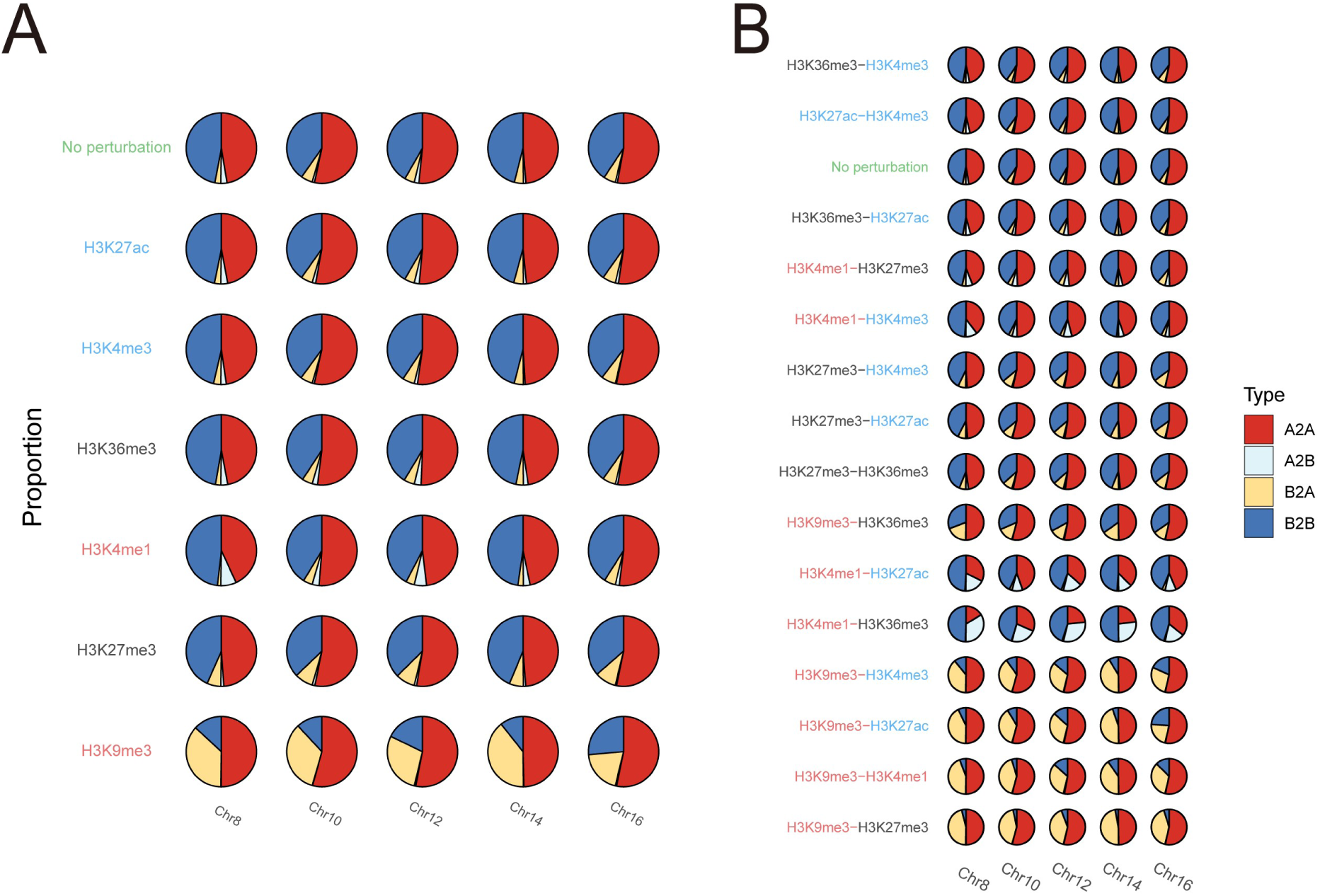
COCOA predicts the compartment patterns for epigenomic perturbation experiments. The row where green “No perturbation” (Reference) indicates the comparison between the predicted CM and ground truth for unperturbed data. Red, black, and blue fonts in the vertical axis labels indicate high, medium, and low impact of the perturbed epigenomic signal on the chromatin compartment patterns, respectively. **(A)** Proportion of the compartment pattern matching between the predicted CM and the ground-truth CM from the 1-perturbation epigenomic combinations. **(B)** Proportion of the compartment pattern matching between the predicted CM and the ground-truth CM from the 2-perturbation epigenomic combinations.

To further analyse the contribution of individual epigenomic signals to the maintenance of chromatin compartment patterns, we conducted keep-one epigenomic signal perturbation (keep-one perturbation) experiments. In these experiments, we perturbed the epigenomic data by maintaining the selected epigenomic signal data while setting all other epigenomic signal data to their minimum values. Subsequently, we utilized COCOA to predict the chromatin compartment patterns for the perturbed data. The keep-one perturbation experiments revealed that the predicted CM from H3K9me3 or H3K4me1 signal partially overlapped with the experimental CM for the unperturbed data, while those predicted CMs from other epigenomic signals exhibited distinct differences from the experimental CM (Figure S4A and Figure S4B). This result reinforced the importance of H3K9me3 and H3K4me1 for predicting the status of chromatin architecture.

We next investigated the effects of two epigenomic signals on the chromatin compartmentalization through two epigenomic signal perturbation (two-perturbations) experiments. We observed that H3K9me3 and H3K4me1 signals play dominant roles in determining A/B compartment patterns. When H3K9me3 or H3K4me1 signal was perturbed along with H3K27ac, H3K27me3, H3K4me1 or H3K4me3 signal, the predicted chromatin compartment patterns exhibited significant changes compared to those for the unperturbed data (Figure S3B). Notably, perturbing the H3K9me3 signal gave rise to compartment B-to-A changes, while perturbing the H3K4me1 signal resulted in compartment A-to-B changes (Figure 4B). Simultaneous perturbations of H3K9me3 and H3K4me1 signal exhibited the greatest impact on the changes in chromatin compartment patterns among all the two-perturbations experiments.

Taken together, COCOA facilitates investigations into the role of epigenomic signals in determining chromatin compartmentalization through *in silico* epigenomic perturbation experiments. Our results suggest that H3K9me3 and H3K4me1 signals are crucial for maintaining the chromatin compartment patterns.

### COCOA shows robust performance of model predictions at different resolutions

To evaluate the performance of COCOA at different resolutions, we used the model trained at 25kb to predict the fine-scale CM using resolution-specific inputs. We first used the trained model to predict 10kb CM for chromosome 16, 17 and 18 data sets and evaluated the performance of the model predictions. The results shown in Table S4 indicate that COCOA achieves consistent and competitive scores on all three test sets. We further evaluated the correlations between the predicted compartment patterns and the ground truth patterns. The predicted 10 kb CM is highly correlated with and the experimental CM (Figure 5A). A similar high correlation was observed between the predicted compartment states and the ground truth states (PC1 of CM). The compartment misclassification rate indicates that COCOA generates CMs containing reliable chromatin compartment information (Figure S5A). The predicted CM exhibits similar plaid patterns to the experimental CM, which correspond well with the epigenomic signals (Figure 5B and Figure 5C). In addition, the modularity of the predicted CM aligns with that of the experimental CM on the whole (Figure S5C). We also observed consistent and significant shifts in epigenomic modification signals within 150kb neighbourhoods around A/B compartment boundaries, in both predicted and experimental CM, which are distinguishable from randomly selected genomic loci (Figure 5D).

**Figure 5.**
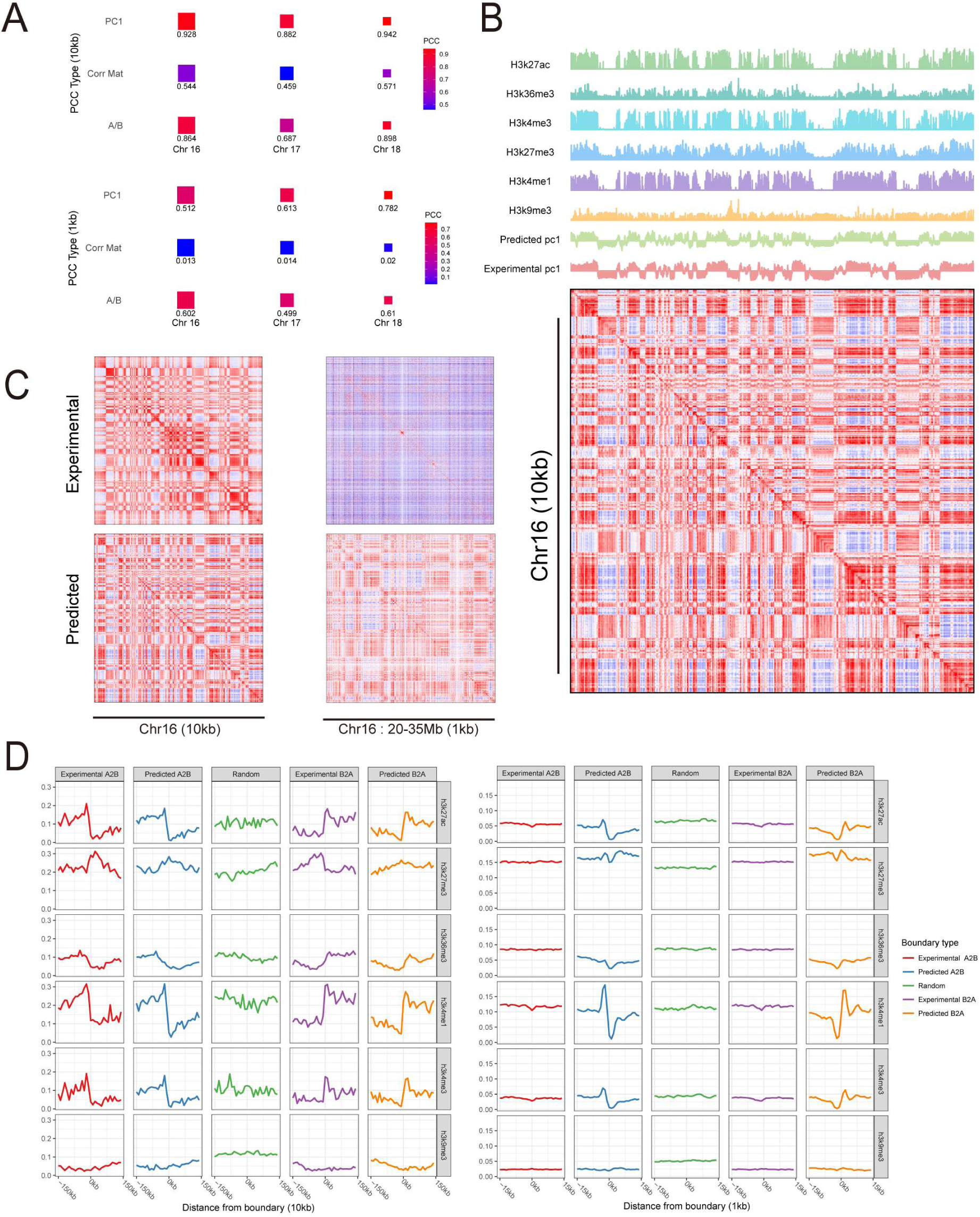
Predicting resolution-specific compartment pattern with COCOA. **(A)** Correlation coefficient between the predicted CM and the experimental CM at 10kb and 1kb resolutions. The rows labeled ‘PC1’, ‘Corr Mat’ and ‘A/B’ represent the correlation of the PC1 of the CMs, the correlation between the predicted CM and the experimental CM, and the correlation of the binarized PC1s, respectively. **(B)** The predicted CM accurately capture the histone modification signal waveforms for chr16 at 10kb resolution. **(C)** Typical region of the predicted CM and the experimental CM at 10kb and 1kb resolutions. At 1kb resolution, the experimental CM exhibits high noise levels and lacks a recognizable plaid pattern, while the predicted CM demonstrates a clear plaid pattern **(D)** Shifts observed in modification signals around compartment boundaries in both predicted and experimental CMs at 10 kb and 1 kb resolutions. At 10 kb resolution, both predicted and experimental CMs display meaningful shifts, whereas at 1 kb resolution, the experimental CM approaches random results, while the predicted CM still shows significant biological shifts.

To evaluate COCOA’s performance at ultra-high resolution, we employ it to predict 1kb for chromosome 16, 17 and 18, using the model trained at 25kb resolution. Similar to the evaluation at 10kb resolution, we assessed the performance metrics and correlations. The results show that COCOA achieves robust performance across a wide range of scores, but obtains scores close to 0 for the CM correlation coefficient (Figure 5A, Table S4). This may be attributed to the sparsity of the deeply-sequenced experimental CM at ultra-high resolution (i.e., ∼2.6–4.5 billion uniquely mapped reads with ∼150X coverage per nucleosome) [47], which is challenging to define as the ground truth. As the CM size increases, the mean error evaluation narrows the gap, producing similar scores. However, using PCA-based correlation or compartment misclassification rates, we can partially mitigate the sparsity issue and obtain reliable scores (Figure S5B). Therefore, we visualized the experimental and predicted CM by heatmaps (Figure 5C). We found that the experimental CM shows vaguely visible plaid patterns and is filled with noise-induced thin lines. In contrast, the predicted CM remains consistent with these fuzzy patterns but displays more apparent compartmentalization patterns (Figure 5C and Figure S5D). Moreover, we investigated the shifts of epigenomic modification signals between the predicted CM and the experimental CM. Surprisingly, the shifts observed in the experimental CM are similar to those in randomly selected genome loci, while the predicted CM shows significant and biologically meaningful shifts (Figure 5D).

### COCOA accurately predicts cell-type-specific chromatin compartment patterns

Because epigenomic data are cell-type-specific, we tested whether COCOA can accurately predict chromatin compartment patterns in different cell types. We first applied COCOA to GM12878 datasets and generated the predicted CM for multiple chromosomes. The experimental CM obtained from the Hi-C data of GM12878 [6] served as the ground truth for comparison. The results indicated that high correlations were observed between the two CMs and between the two principal components (PC1s) of the CMs (Figure 6A and Figure S6A). Figure 6B revealed that the compartment misclassification rates were all below 20%. Furthermore, the predicted CM presents the plaid patterns consistent with the experimental CM, achieving stable and competitive scores in terms of both error and the image similarity (Figure 6C and Table S5). Taken together, these data suggest that COCOA reliably predicts the cell-type-specific chromatin compartment patterns.

**Figure 6.**
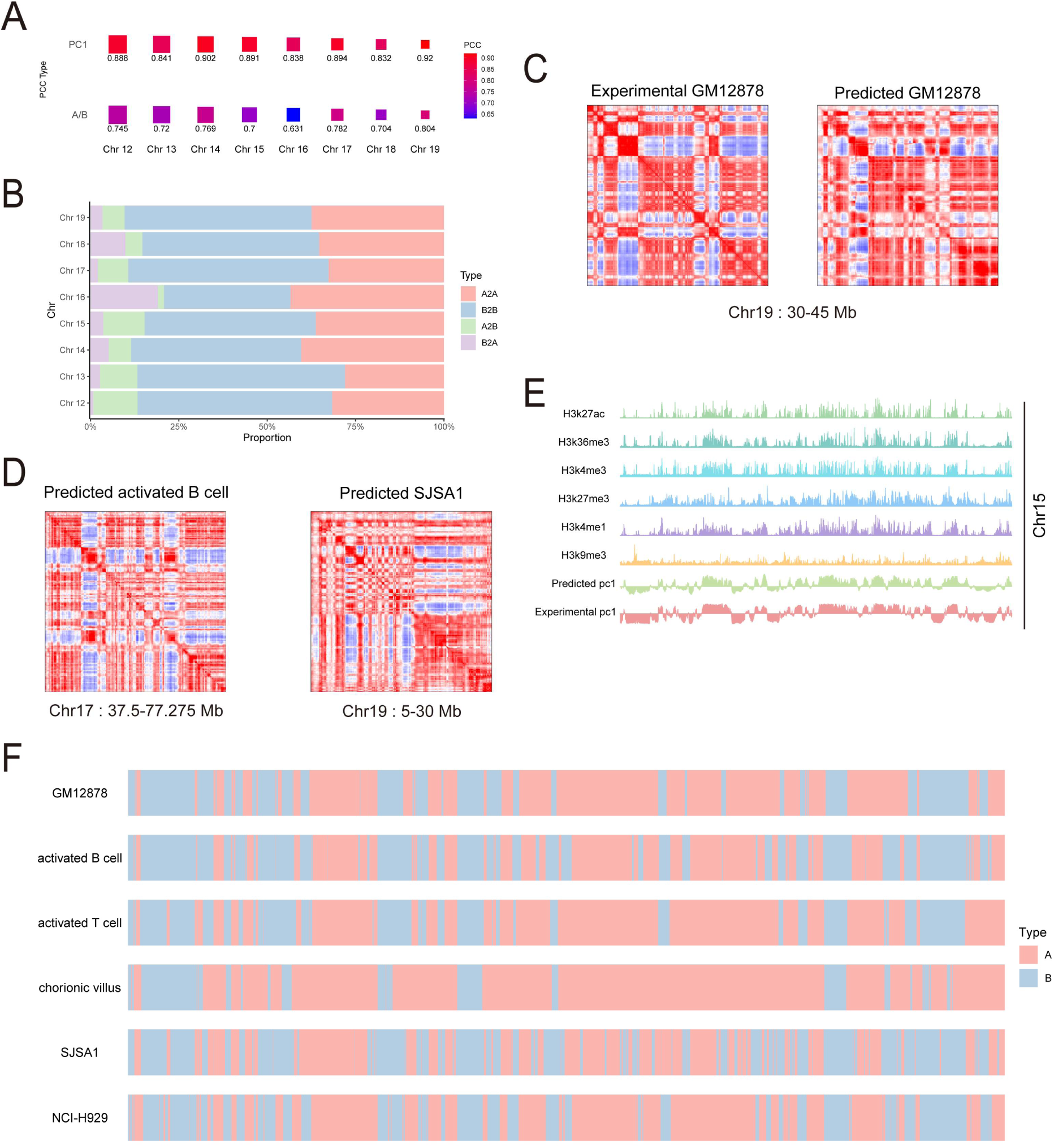
Cell-type-specific prediction of compartment pattern by COCOA. **(A)** Correlation coefficient between the predicted CM and the experimental CM on GM12878 datasets. The rows ‘PC1’ and ‘A/B’ represent the correlation of CM’s PC1s and the correlation of binarized PC1s, respectively. **(B)** Proportion of matching compartment patterns between the predicted CM and the experimental CM on GM12878 datasets. **(C)** Typical regions showing the predicted CM and the experimental CM patterns on GM12878 datasets. **(D)** Example regions illustrating the predicted CM patterns on SJSA1 and activated B cells datasets. **(E)** Precisely matching of predicted CM patterns with the waveform of histone modification signals on chr15 in GM12878 datasets. **(F)** Systematic comparison of chromatin compartment status on chr15 across GM12878, activated B cells, activated T cells, chorionic villus, SJSA1 and NCI-H929 datasets.

Having established COCOA’s capability to predict CMs across diverse cell types, we proceeded to predict CMs in five additional cell types encompassing tissues, diseases, and primary cells where chromatin conformation had not been sequenced. The predicted CM for the five data sets displayed obvious plaid patterns (Figure 6D and Figure S6B). The predicted A/B compartments in SJSA1, NCI-H929, and activated B cells were clearly defined and noise-free, while the predicted CMs of activated T cells and chorionic villus tissues displayed slightly diminished performance. To gain insight into the effects of chromatin compartmentalization in disease and differentiation, we systematically compared the patterns across different cell types. Using predicted CM from GM12878 data as a benchmark, we examined commonalities and differences in chromatin region patterns based on histone modification track information (Figure 6E). The results revealed similar patterns between GM12878 and most other cells, albeit with variations in certain regions (Figure 6F). Notably, activated B cells, being immune cells akin to GM12878, exhibited a comparable compartment pattern. Similarly, activated T cells demonstrated an analogous pattern. In contrast, SJSA1, NCI-H929, and chorionic villus tissues exhibited distinct compartment patterns.

## Discussion

In this study, we developed a deep neural network framework called COCOA, which incorporates six types of epigenomic modification signals to accurately predict fine-scale resolution chromatin compartment patterns. These epigenomic signals data are readily accessible in ENCODE database [53] for various cell lines, in vitro differentiated cells, primary cells, and tissues. To process the raw epigenomic data, we employed resolution-specific pre-processing to bin the data into mated inputs from different genomic positions. COCOA then uses the bidirectional feature reconstruction module to extract track features from these mated inputs, then fuses these track features to contact features using the cross attention fusion module. Eventually, these contact features are converted to chromatin compartment patterns by the residual feature reduction module. COCOA predict directly long-range chromatin compartment patterns without considering short-range interactions [28,29,35]. Our results demonstrate that COCOA accurately predicts the same chromatin compartment patterns as the experimental CM, with consistent epigenomic signals shifts of these patterns (Figure 2D). During model evaluation, COCOA achieves excellent performance with robust reproducibility scores on the test sets. Furthermore, the predicted CM and its PC1 show a high correlation with the experimental CM and PC1. The compartment misclassification rates of the predicted CM remain below 10% and are independent of chromosome length.

With COCOA’s accurate prediction of chromatin compartmentalization, it becomes possible to perform *in silico* epigenomic perturbation to study the influence of histone modification signals on chromatin compartmentalization. By generating predicted CM using different perturbed epigenomic data, we found that H3K9me3 has strong impact on chromatin compartment patterns, followed by H3K4me1. In contrast, H3K27me3 and H3K36me3 have a moderate level of impact, and H3K27ac and H3K4me3 have low impact. Interestingly, COCOA predicted that perturbation of H3K9me3 signal led to compartment B-to-A changes, while perturbation of H3K4me1 signal resulted in compartment A-to-B switches. Additionally, H3K9me3 and H3K4me1signals play dominant roles in determining chromatin compartment patterns when they are perturbed together with other epigenomics signals in two-perturbations experiments.

Furthermore, we explored the performance of COCOA’s predictions across different resolutions and cell types. For prediction at 10kb resolution, COCOA exhibited the same outstanding performance as predicted at the trained resolution (25kb). Recognizing the significance of high resolution in chromatin interaction data analysis, we investigated whether COCOA can make good prediction at 1kb resolution. Unfortunately, even with a high sequenced depth [47], the experimental CM at 1kb resolution contains excessive noise lines and barely discernible plaid pattern. Therefore, we analysed histone modification shifts at compartment boundaries and mapped heatmaps of the predicted CM at different genome ranges. Surprisingly, the predicted CM displayed clearer plaid patterns and exhibited more biologically meaningful shifts compared to the experimental CM and randomly selected loci. We then evaluated the performance of COCOA in predicting cell-type-specific compartment patterns. Using validated Hi-C data of GM12878, our results demonstrated that COCOA can correctly infer chromatin compartment patterns from epigenomic data on unseen cell lines.

While this work presents promising results, it also has several potential areas for improvements. Firstly, as a data-driven approach, COCOA relies on moderately good quality training sets to achieve high performance by incorporating potential information from bidirectional epigenomic data. In addition, we observed that the transfer capacity of COCOA in cross cell lines experiments is affected by the epigenomic data quality. Developing new data processing schemes may prove beneficial in solving this issue. Secondly, in challenging task such as high-volume fine-scale resolution CM prediction and *in silico* epigenomic signal perturbation experiments, COCOA requires significant run time and substantial computational resources. To alleviate this computational burden, parallel CM generation and distributed implementations can be explored as feasible approaches [55]. Thirdly, we also preliminarily explored the influences of histone modification signals on A/B chromatin compartmentalization in HFF datasets by *in silico* epigenomic perturbation experiments. However, more systematically studying the combined impacts of epigenomic modifications in relation to complex chromatin compartmentalization on different cell lines would benefit from further experimental evidence. Lastly, COCOA’s predictions for fine-scale chromatin compartmentalization information in diseases, tissues and primary cells have not been thoroughly explored. In future, it would be interesting to explore the impact of the chromatin compartment alteration on cell differentiation and disease occurrence by integrating epigenomics data with other omics and phenotypic data.

## Code availability

The source codes were implemented in Python and can be freely accessed on GitHub at the following link: https://github.com/onlybugs/COCOA

## Data availability

The datasets used in this study are listed in Table S1, S2 and can be obtained from the ENCODE project and the 4D Nucleome Data Portal.

## CRediT author statement

**Kai Li:** Conceptualization, Methodology, Software, Data curation, Visualization, Formal analysis, Writing - original draft. **Ping Zhang:** Conceptualization, Writing - review. **Jinsheng Xu:** Data curation. **Zi Wen:** Data curation. **Junying Zhang:** Data curation. **Zhike Zi:** Writing - review & editing, Resources, Conceptualization. **Li Li:** Writing - review & editing, Project administration, Resources. All authors have read and approved the final manuscript.

## Acknowledgments

This work was supported by the Huazhong Agricultural University Scientific and Technological Self-innovation Foundation (to L.L.), Numerical computations were performed on the Hefei Advanced Computing Center. Z.Z. would like to acknowledge the support from Guangdong Provincial Key Laboratory of Synthetic Genomics and Shenzhen Key Laboratory of Synthetic Genomics (ZDSYS201802061806209). We thank Li lab members for providing feedback on the earlier version of the manuscript.

