## Supplemental materials for "COCOA: A Framework for Fine-scale Mapping Cell-type-specific Chromatin Compartmentalization Using Epigenomic Information"

### Note S1

**Model training and hyper-parameters details.** All the training and validation processes were conducted on NVIDIA 1080 GPU and 300 GB of memory. During evaluation, we performed prediction and combination on Intel(R) Xeon(R) CPU E5–2696 v4 with 503 GB of memory. The model is trained with a batch size of 16 for 120 epochs and adopts the Adam optimizer with an initial learning rate of 5e-4 (lr_init=5e-4). In addition, we employed early stopping to prevent overfitting and used the step learning rate scheduler (lr_n= lr_init*dr^(n/10), where n denotes the current epoch, dr is decay ratio).

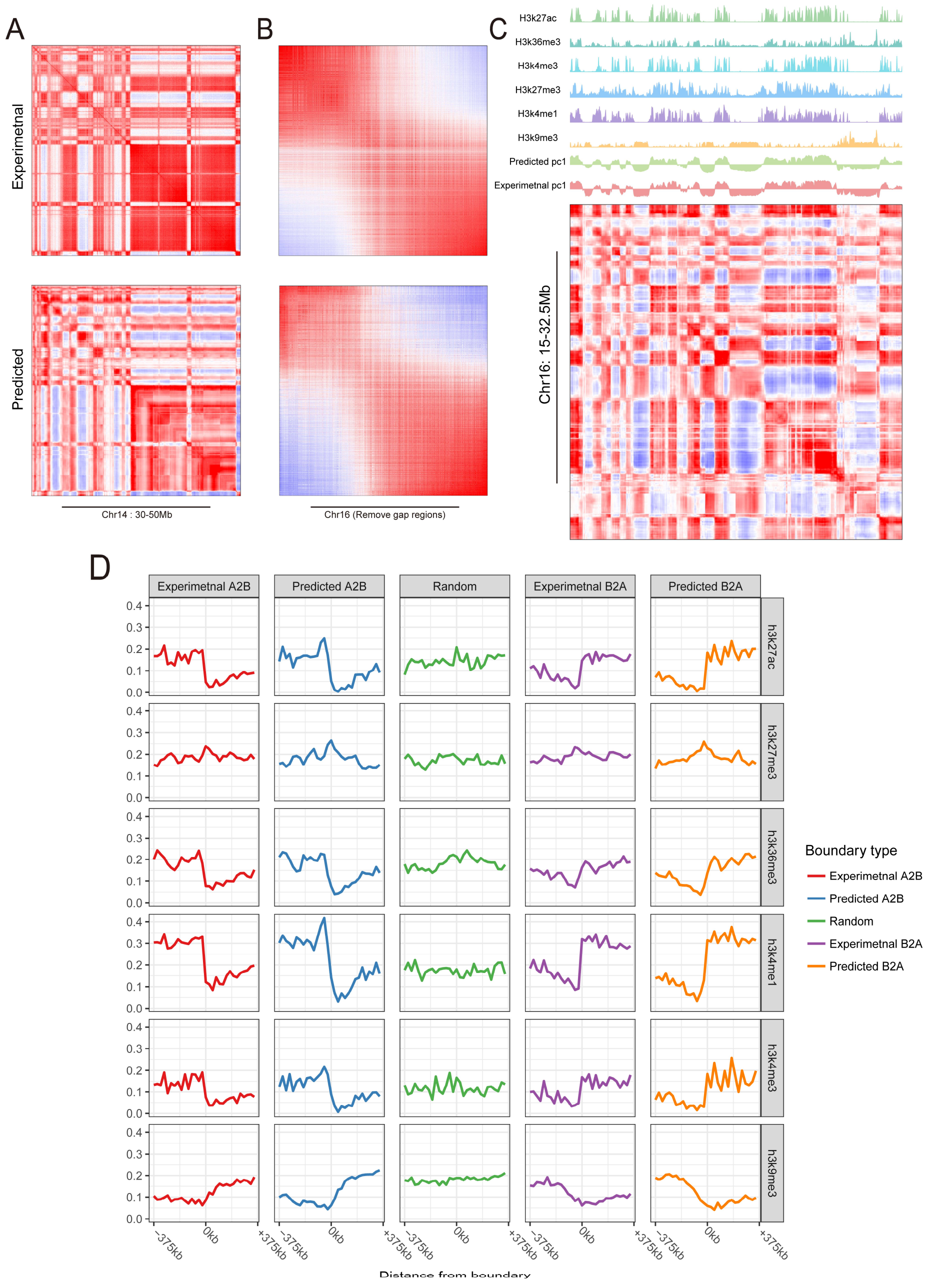


**Figure S1 COCOA precisely infers biological compartment patterns from epigenomic data**

**(A)** Representative region illustrating predicted and experimental CM on Chr14: 30-50Mb.

**(B)** Heatmaps of the experimental CM and the predicted CM, sorted according to their respective PC1 sizes.

**(C)** The predicted CM exhibits patters that align precisely with the waveform of histone modification signal.

**(D)** Representative shifts in modification signals within 375 kb neighbourhoods surrounding compartment boundaries in both predicted and experimental CMs.

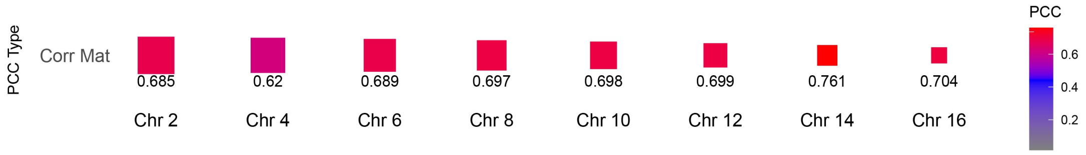


**Figure S2 Correlation coefficient between the predicted CM and the experimental CM**

Correlation coefficient between the predicted CM and experimental CM. The "Corr Mat" row represents the average correlation coefficient of each column of the CM.

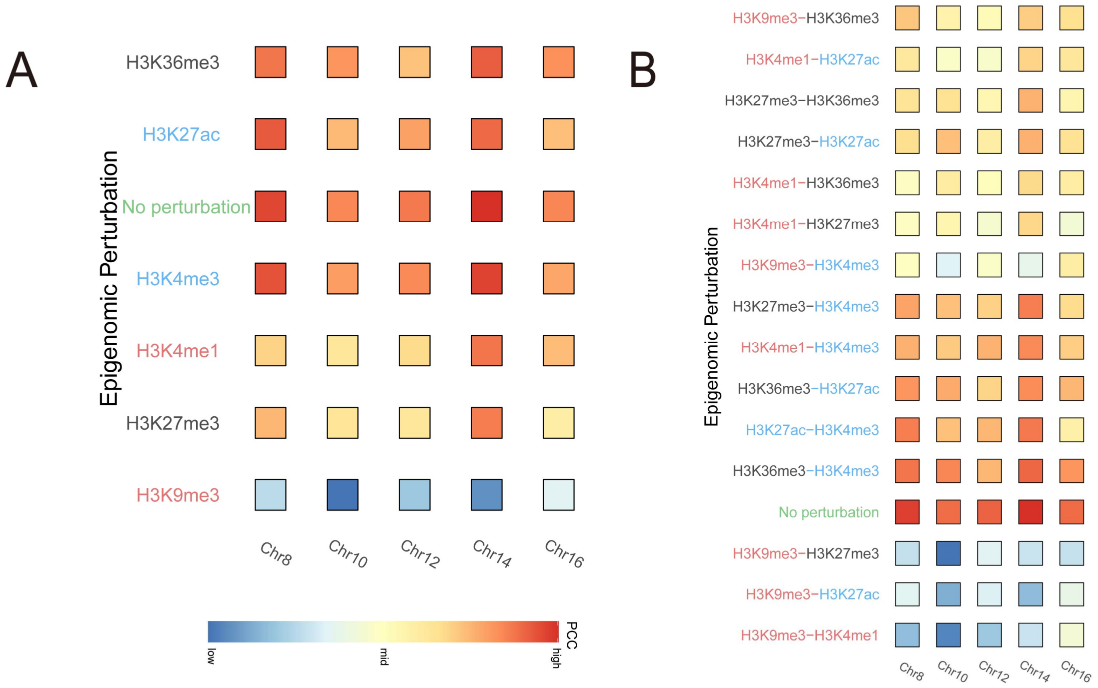


**Figure S3 COCOA performs the results of the epigenomic one-perturbation and the two-perturbation experiments**

The row where green "No perturbation" (Reference) indicates the comparison between the predicted CM and ground truth for unperturbed data. Red, black, and blue fonts in the vertical axis labels indicate high, medium, and low impact of the perturbed epigenomic signal on the chromatin compartment patterns, respectively.

**(A)** The average correlation coefficients between the predicted CM from one-perturbation experiments and the experimental CM from the unperturbed data. The corresponding PC1 correlation coefficients are also shown for different test chromosomes.

**(B)** The similar PC1 correlation coefficients from two-perturbation experiments.

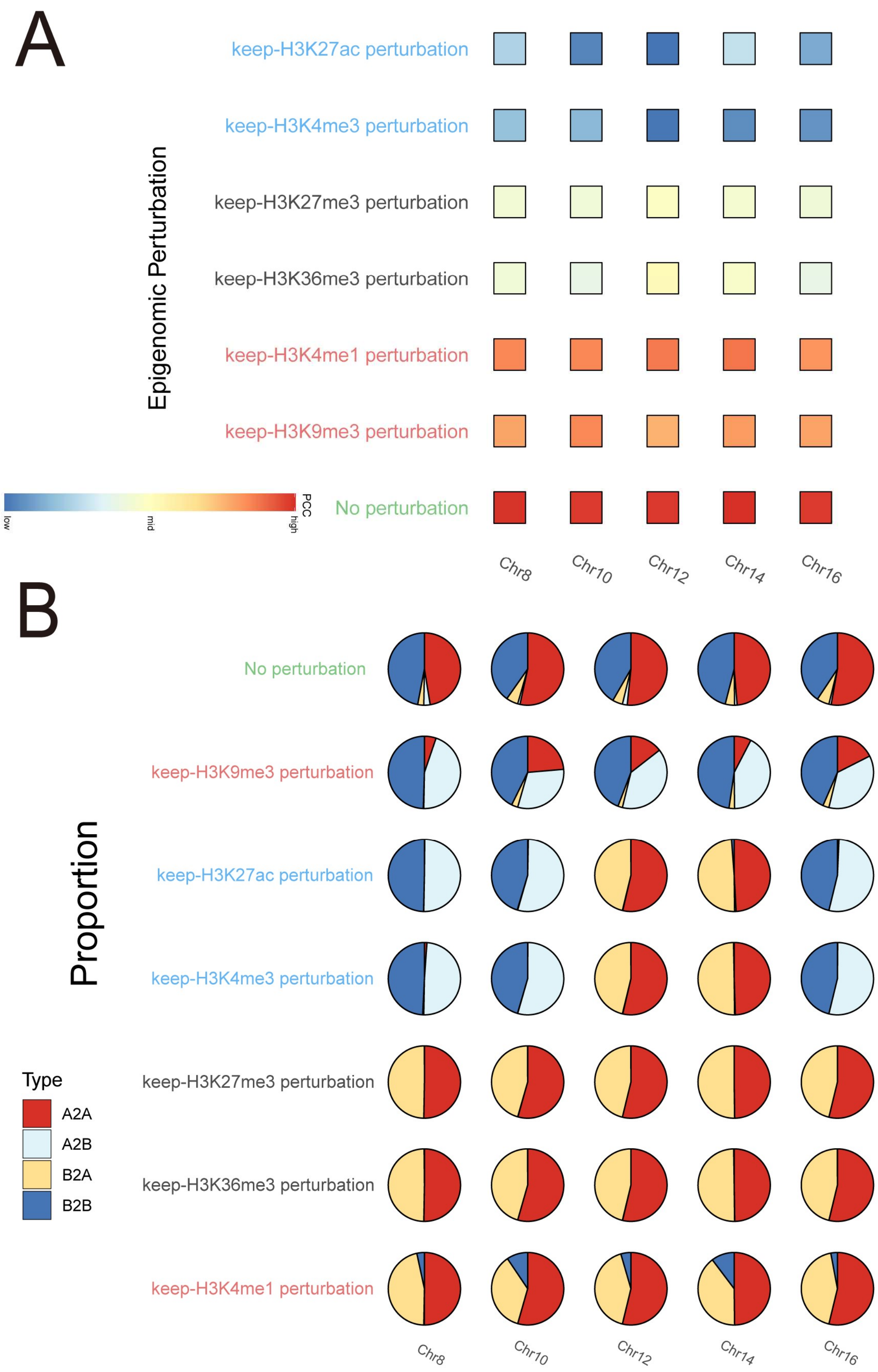


**Figure S4 COCOA performs the results of the keep-one perturbation experiments**

**(A)** The average correlation coefficients between the predicted CM from keep-one perturbation experiments and the experimental CM from the unperturbed data. The corresponding PC1 correlation coefficients are also shown for different test chromosomes.

**(B)** Proportion of the compartment pattern matching between the predicted CM and the ground-truth CM from the keep-one perturbation experiments.

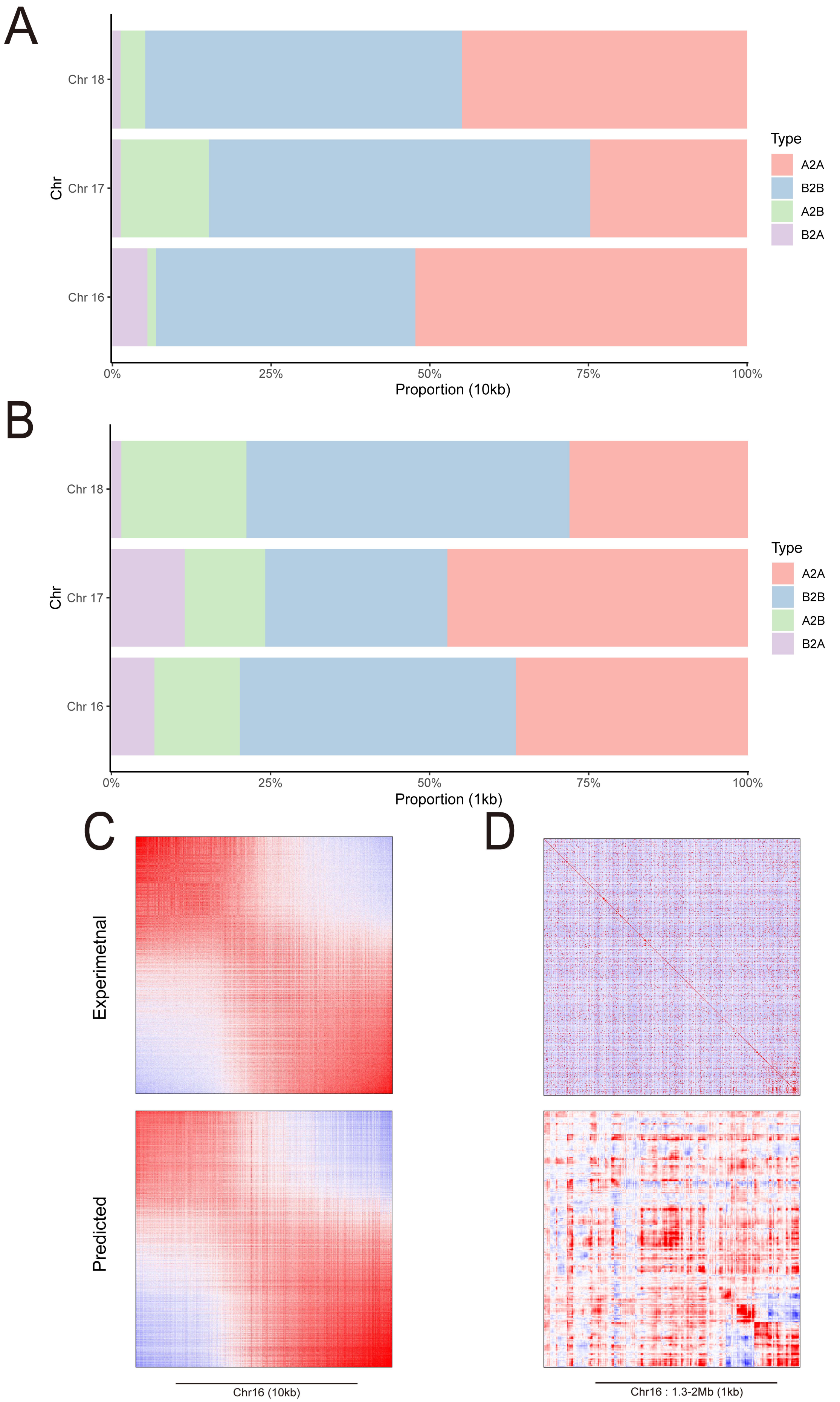
A
Chr 18
Chr 17
Chr 16
Chr
0%
25%
50%
75%
100%
Proportion (10kb)
Type
A2A
B2B
A2B
B2A
B
Chr 18
Chr 17
Chr 16
Chr
0%
25%
50%
75%
100%
Proportion (1kb)
Type
A2A
B2B
A2B
B2A
C
D
Experimetnal
Predicted
Chr16 (10kb)
Chr16 : 1.3-2Mb (1kb)

**Figure S5 COCOA achieves reliable performance across different resolutions**

**(A)** Proportion of compartment pattern matching between the predicted CM and the experimental CM at 10kb resolution. The red and blue bars represent the proportion of compartments that overlap between the predicted CM and experimental CM. The green and purple bars indicate the proportion of compartments that differ between the predicted CM and experimental CM.

**(B)** Proportion of compartment pattern matching between the predicted CM and the experimental CM at 1kb resolution.

**(C)** Heatmaps of the experimental CM and the predicted CM, sorted according to their respective PC1 sizes at 10kb resolution.

**(D)** Typical region of the predicted CM and the experimental CM at 1kb resolutions.

A

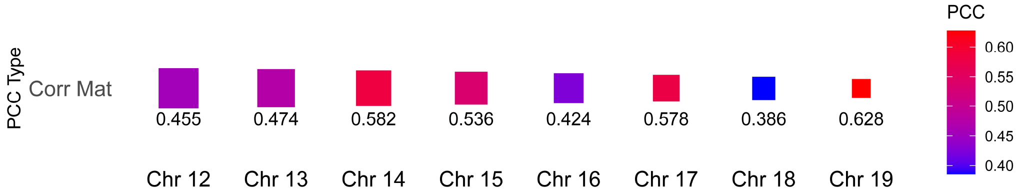


B

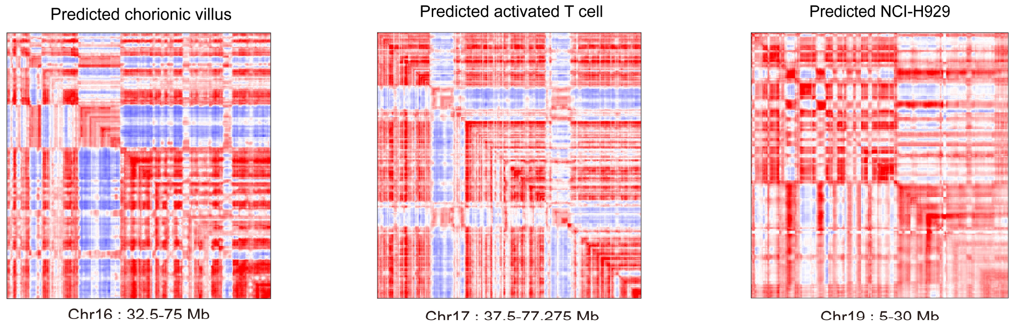


**Figure S6 COCOA enables reliable cell-type-specific prediction of compartment pattern**

**(A)** Correlation coefficient between the predicted CM and experimental CM on GM12878 datasets. The “Corr Mat” row represents the average correlation coefficient of each column of the CM.

**(B)** Example regions illustrating the predicted CM patterns on chorionic villus, activated T cell and NCI-H929 datasets

**Table S1 Micro-C and Hi-C data**

| Type | Data | Accession number |
|---|---|---|
| Micro-C | HFFc6 | 4DNESWST3UBH |
| Hi-C | GM12878 | 4DNFIXP4QG5B |

**Table S2 ChIP-seq data**

| Type | HFFc6 | GM12878 | Activated T cell | Activated B cell | NCI-H929 | SJSA1 | Chorionic villus |
|---|---|---|---|---|---|---|---|
| H3K4me3 | ENCFF598MLG | ENCFF012DMX | ENCFF641ONK | ENCFF399LTS | ENCFF754JLX | ENCFF596HJB | ENCFF548VTO |
| H3K27ac | ENCFF644GJB | ENCFF798KYP | ENCFF169DFQ | ENCFF490TAG | ENCFF612UYR | ENCFF005KFY | ENCFF884SVO |
| H3K27me3 | ENCFF854VTY | ENCFF677PYB | ENCFF328DSH | ENCFF394DBD | ENCFF588PCB | ENCFF548QPW | ENCFF840RKG |
| H3K4me1 | ENCFF834VWT | ENCFF190RZM | ENCFF036KBQ | ENCFF110AUF | ENCFF289DWP | ENCFF574PIN | ENCFF635XBE |
| H3K36me3 | ENCFF431XVV | ENCFF345QSP | ENCFF247PKZ | ENCFF942SCO | ENCFF863VMA | ENCFF532OYN | ENCFF259MAY |
| H3K9me3 | ENCFF542CZT | ENCFF701GHA | ENCFF071KRK | ENCFF710FWU | ENCFF426TFY | ENCFF059AXW | ENCFF409PTA |

The contents of the table are the Accession number of the corresponding data in the ENCODE database.

**Table S3 Summary table (performance evaluation)**

| Chr | MAE | SSIM | PSNR |
|---|---|---|---|
| 2 | 0.2318 | 0.3500 | 11.94 |
| 4 | 0.2667 | 0.3086 | 10.81 |
| 6 | 0.1731 | 0.4220 | 13.84 |
| 8 | 0.2216 | 0.4261 | 12.19 |
| 10 | 0.1971 | 0.4321 | 12.76 |
| 12 | 0.1916 | 0.4308 | 13.18 |
| 14 | 0.2104 | 0.4394 | 12.16 |
| 16 | 0.1601 | 0.4352 | 13.82 |

**Table S4 Summary table (multiple resolution)**

| Chr | MAE | MSE | SSIM | PSNR |
|---|---|---|---|---|
| 16 (10k) | 0.2404 | 0.0758 | 0.1750 | 11.20 |
| 17 (10k) | 0.2051 | 0.0565 | 0.1961 | 12.48 |
| 18 (10k) | 0.2169 | 0.0614 | 0.1828 | 12.12 |
| 16 (1k) | 0. 0603 | 0.0123 | - | 19.10 |
| 17 (1k) | 0.1389 | 0.0273 | - | 15.64 |
| 18 (1k) | 0.1634 | 0.0346 | - | 14.61 |

'-' indicates metrics that are not available due to computational resource constraints.

**Table S5 Summary table (cell-type-specific prediction)**

| Chr | MSE | MAE | SSIM | PSNR |
|---|---|---|---|---|
| 12 | 0.0620 | 0.2034 | 0.3566 | 12.08 |
| 13 | 0.0871 | 0.2615 | 0.2690 | 10.60 |

| | | | | |
|---|---|---|---|---|
| 14 | 0.1212 | 0.3000 | 0.3200 | 9.16 |
| 15 | 0.0836 | 0.2466 | 0.3362 | 10.78 |
| 16 | 0.0804 | 0.2342 | 0.3208 | 10.95 |
| 17 | 0.0667 | 0.2125 | 0.3981 | 11.74 |
| 18 | 0.0786 | 0.2376 | 0.2564 | 11.04 |
| 19 | 0.0585 | 0.1953 | 0.4462 | 12.33 |
